# Reassessing Number-Detector Units in Convolutional Neural Networks

**DOI:** 10.64898/2026.03.07.710304

**Authors:** Nhut Truong, Shahryar Noei, Alireza Karami

## Abstract

Convolutional neural networks (CNNs) have become essential models for predicting neural activity and behavior in visual tasks. However, their ability to capture higher-level cognitive functions, such as numerosity discrimination, remains debated. Numerosity, the ability to perceive and estimate the number of items in a visual scene, is often proposed to rely on specialized number-detector units within CNNs, analogous to number-selective neurons observed in the brain. In this study, we use CORnet, a biologically inspired CNN architecture inspired by the organization of the primate visual system. To address a limitation of classical Representational Similarity Analysis (RSA)—its assumption that all units contribute equally—we apply pruning, a feature selection approach that identifies the units most relevant for explaining behavioral similarity structure. Our results show that number-detector units are not critical for population-level representations of numerosity, challenging their proposed role in previous studies.

## 1 Introduction

Early breakthroughs in the study of biological vision served as the foundation for convolutional neural networks (CNNs; Lindsay 2021). Like the brain, these hierarchical models consist of several feedforward layers, with each layer comprising numerous artificial units that mimic neurons. Since then, CNNs have evolved into state-of-the-art models for predicting neural activity and behavior in visual tasks (Khaligh-Razavi and Kriegeskorte 2014; Daniel L. K. Yamins et al. 2014; Daniel L K Yamins and DiCarlo 2016; Cichy et al. 2016). For example, it has been demonstrated that CNNs trained on an object classification task can account for the brain responses of both humans’ and monkeys’ inferior temporal cortex (IT), a key region for object recognition (Khaligh-Razavi and Kriegeskorte 2014).

But what happens when the images contain multiple objects? Perceiving and representing the number of items in a set—known as numerosity—without counting is considered a core and ancient cognitive ability shared by humans and many animal species, often referred to as ‘number sense’ (Dehaene 2011). Specialized neurons, or ‘number neurons’, that are tuned to the number of items in a visual display have been identified in numerically naive monkeys (Viswanathan and Nieder 2013), crows (Wagener et al. 2018), and untrained 10-day-old domestic chicks (Kobylkov et al. 2022), suggesting that numerosity is automatically represented in the brain. Brain imaging studies have also pinpointed regions in the parietal cortex that are responsible for representing numerosity in both adults (Piazza et al. 2004; Castaldi et al. 2019; Karami, Castaldi, Eger, and Piazza 2025) and preverbal infants (Izard, Dehaene-Lambertz, and Dehaene 2008; Hyde et al. 2010; Edwards et al. 2015), demonstrating this ability at the population level. In addition, Karami, Castaldi, Eger, M. Hebart, et al. 2025 combining magnetoencephalography (MEG) with fMRI has shown that numerosity representations emerge rapidly after stimulus onset and evolve over time along the visual hierarchy, from early visual cortex to higher-level associative areas. Additionally, fMRI decoding in the parietal regions of adults has been linked to behavioral number discrimination acuity (Lasne et al. 2019). Collectively, these findings highlight the critical role of parietal brain activity in human number discrimination.

Recently, it has been shown that number-detector units, analogous to number neurons recorded in the prefrontal and parietal cortices of monkeys, can emerge in the final layers of CNNs trained for visual object recognition (Nasr, Viswanathan, and Nieder 2019) and even in completely untrained CNNs (Kim et al. 2021). However, Karami, Truong, and Piazza (2025), using RSA (Kriegeskorte 2008), demonstrated that CNNs fall short of explaining the variance in numerosity representation observed in fMRI data from the human parietal cortex. Further analysis using multidimensional scaling (MDS; Kruskal 1964) revealed significant differences in the geometric structure of numerosity representations between human parietal regions and CNNs (Karami, Truong, and Piazza 2025). In the classical RSA framework used by Karami (2024), all features contribute equally to the final dissimilarity estimate. However, this ‘equal weights’ assumption conflicts with the notion that, when comparing representational dissimilarity matrices (RDMs), certain features may carry more informative content than others. As a result, this approach can underestimate the true correspondence between the model and a specific brain region or behavior (Kaniuth and M. N. Hebart 2022; Tarigopula et al. 2023). Moreover, the classical RSA approach may overemphasize non-relevant units by treating them as equally important as units that carry behaviorally relevant information, such as number-detector units in our case.

To assess the relevance of number-detector units in representing numerosity at the population level within CNNs, we employed a pruning approach. Pruning is a feature selection technique used to identify and retain the most relevant parts of a model, such as specific weights or activations, that best align with the behavior data and improve predictions (Flechas Manrique et al. 2023; Bao and Hasson 2024; Truong, Pesenti, and Hasson 2024). This approach is based on the observation that pretrained models often contain redundant information (Cheng et al. 2015; Frankle and Carbin 2018). Therefore, using the entire model may not be necessary for a specific task, such as numerosity discrimination in our case. Specifically, we pruned different CNN architectures based on the number RDM, which captures the behavioral signature of numerosity perception in humans. This matrix serves as a benchmark for comparing the alignment of CNN representations with human numerosity processing. Our results revealed that number-detector units are not critical for representing numerosity at the population level within these networks. This finding suggests that, while number-detector units may emerge in specific layers of CNNs, they do not play a significant role in capturing the broader, population-level representation of numerosity, as reflected in human behavioral data.

## 2 Methods and experimental setup

### 2.1 Stimuli and Training the CNN

To investigate number-detector units in CNNs we used CORnet-Z and CORnet-S, models with four anatomically mapped areas (V1, V2, V4, and IT) followed by a decoder layer. CORnet-Z is the simplest network in the CORnet family and a lightweight alternative to AlexNet. CORnet-S also has recurrent connectivity and is designed to maximize Brain-Score (Schrimpf et al. 2018). Each anatomically mapped area in the CORnet consists of a single convolution, followed by a ReLU nonlinearity, max pooling and the decoder is a 1000-way linear classifier (Kubilius, Schrimpf, Nayebi, et al. 2018; Kubilius, Schrimpf, Kar, et al. 2019).

We chose CORnet-Z and CORnet-S because it balanced the resemblance to the architectures used by previous studies on numerosity and because it well fit the visual system. We used three versions of the CORnet:

1. the completely untrained version with randomly initialized weights to reveal the effect of architecture alone (Cichy et al. 2016),
2. a version trained on object recognition using the ImageNet dataset (Deng et al. 2009), which contained 1.2 million images of objects over 1000 categories, as it has been used in a previous study by Nasr, Viswanathan, and Nieder (2019),
3. and a version of the network was specifically trained to discriminate between ten numerosity values: 6, 7, 9, 10, 12, 14, 17, 20, 24, and 29. We specifically trained the networks to discriminate between numbers because Mistry et al. (2023) demonstrated that training a CNN for numerosity discrimination significantly reorganizes the number-detector units. To avoid flawed stimulus design (Park 2022), where low-level visual features like size or dot density correlate with the number of dots, we used the method introduced by DeWind et al. (2015) to generate the dot sets. A sample of the generated stimuli used for training the network is shown in Figure 1A. Following the approach of Mistry et al. (2023), we first initialized the network with weights pre-trained on ImageNet, then trained it on the numerosity task for 100 epochs using the Stochastic Gradient Descent (SGD) optimizer with default PyTorch parameters. The code used to train the networks, along with a link to the trained network weights, will be available in the GitHub repository associated with this paper after the anonymous review process.

**Figure 1.**
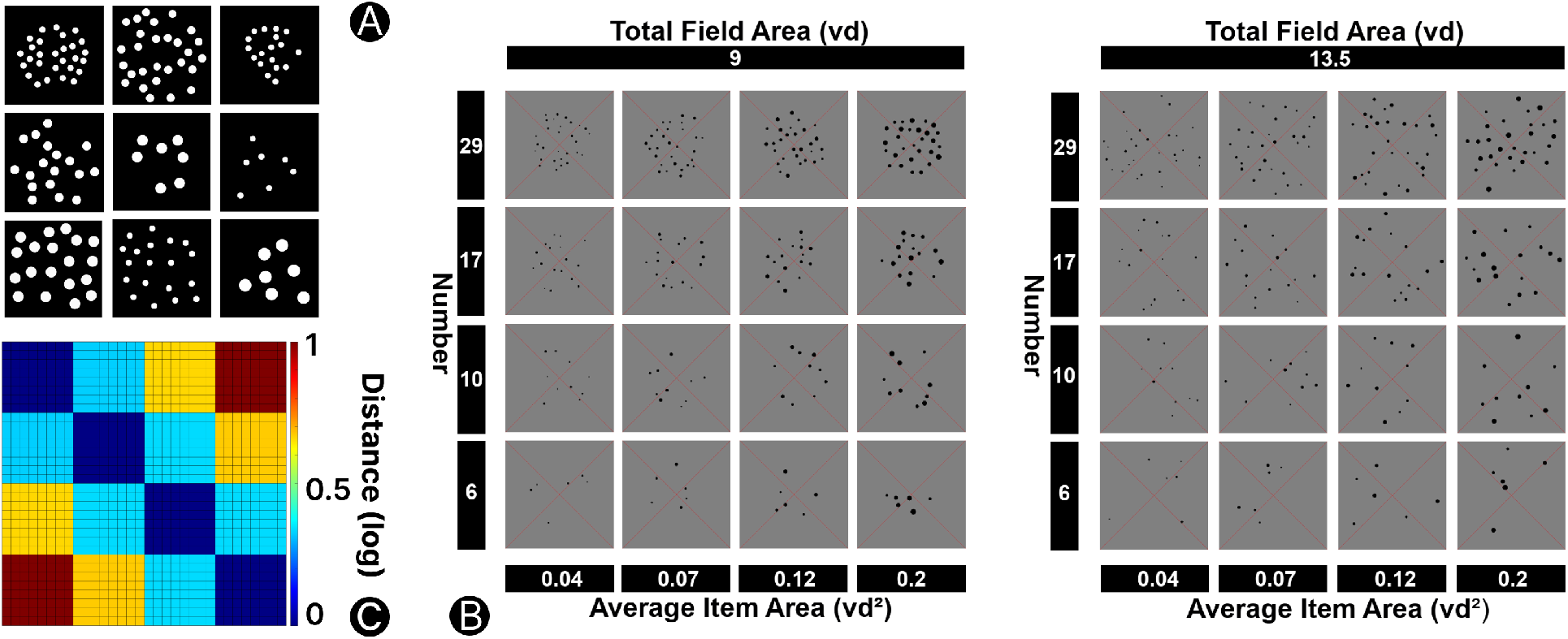
(A) Sample stimuli used for training the networks. (B) Sample stimuli used for testing the networks. (C) Number RDM, reflecting human behavioral data, used to prune the network layers.

After training, the three versions of the model were tested with visual dot sets, where both the number of dots and low-level visual features (average item area and total field area) varied across 32 different conditions: 4 numerosities (6, 10, 17, 29), 4 average item areas, and 2 total field areas (Figure 1B). Each input image was 500 × 500 pixels. We selected four layers of the network-analogous to visual brain areas (V1, V2, V4, and IT) and extracted the activations of all nodes in each layer. The results from 100 instances of each condition were averaged to produce a single activity vector for each condition from the output of each layer.

### 2.2 Identification of Number-Detector Units

Following Nasr, Viswanathan, and Nieder (2019) we used ANOVA to find number-detector units. Specifically, a three-way analysis of variance (ANOVA) with three factors - numerosity, total field area, and average item area - was applied to select number-detector units. The goal was to identify units that had a significant change in response across different numerosities while maintaining an invariant response across variations in total field area and average item area, as well as across the three interactions among pairs of factors, and the interaction among all of the three. A unit is marked as a number-detector if it shows a significant change for numerosity (p < 0.01) but no significant change for the other two factors or any of the interactions. Units that did not meet these criteria were classified as non-selective. This method of selecting number-detector units is analogous to the method that has been used to detect numerosity-sensitive neurons in monkeys and humans.

### 2.3 Representational Similarity Analysis between Number RDM and Network RDM

To create the CNNs’ RDMs, we used the 32 activity vectors obtained by averaging the 100 instances per condition. We chose 100 instances to address concerns about the limited number of sample images used in previous studies, such as Nasr, Viswanathan, and Nieder (2019), which were criticized for this limitation (Zhang and Wu 2020). The CNNs’ RDMs were constructed using 1 – Pearson correlation between the activations of each layer for each pair of conditions. The number RDM (Figure 1C) was based on the logarithmic distance between the pairs of conditions in terms of numerosity. We then compute the correlation between each CNN’s RDM and the number RDM.

#### 2.3.1 Pruning the Layers of Models

The pruning algorithm, which is adapted from Tarigopula et al. (2023), involves three steps. First, the importance of each unit is assessed by removing it from the full set of units. Each time a unit is removed, a new RDM is computed, and its score is compared to the number RDM. A significant drop in the score compared to the full set RDM indicates that the unit is important for matching the number RDM, while a smaller drop or an increase in score suggests the unit is unimportant or possibly encoding noise. Second, all units are ranked based on their importance scores, from highest to lowest. Third, starting with an empty activation vector, units are sequentially added back in the order of their ranking. After each addition, the fit between the RDM derived from the new embedding and the number RDM is re-evaluated. We truncate and select the set of neurons when the highest RSA score is achieved, and refer these units as the ‘retained units’ after pruning.

## 3 Results

### 3.1 Retained Units After Pruning and Number-Detector Units are Distinct

Table 1 presents the number of units selected by two methods: pruning and ANOVA. In both models, the number of retained units after pruning is significantly higher than the number of number-detector units identified by ANOVA. ANOVA also results in significant more units in CORnet-S compared to Z. Moreover, pruning removed the largest proportion of units in the IT layer, while the ANOVA method showed no significant differences across layers. There is no noticeable difference between the ImageNet-trained and DeWind-trained models. Interestingly, both retained units and number-detector units are found in untrained networks, consistent with the findings of Kim et al. (2021). Regarding the overlap between the units selected by the two methods - defined as len(A ∩ B) / min(len(A), len(B)) - the overlap remains below 0.05 in most cases, except for CORnet-S DeWind (0.53) and CORnet-Z ImageNet (0.15). This indicates that the two methods generally select distinct sets of units.

**Table 1:**
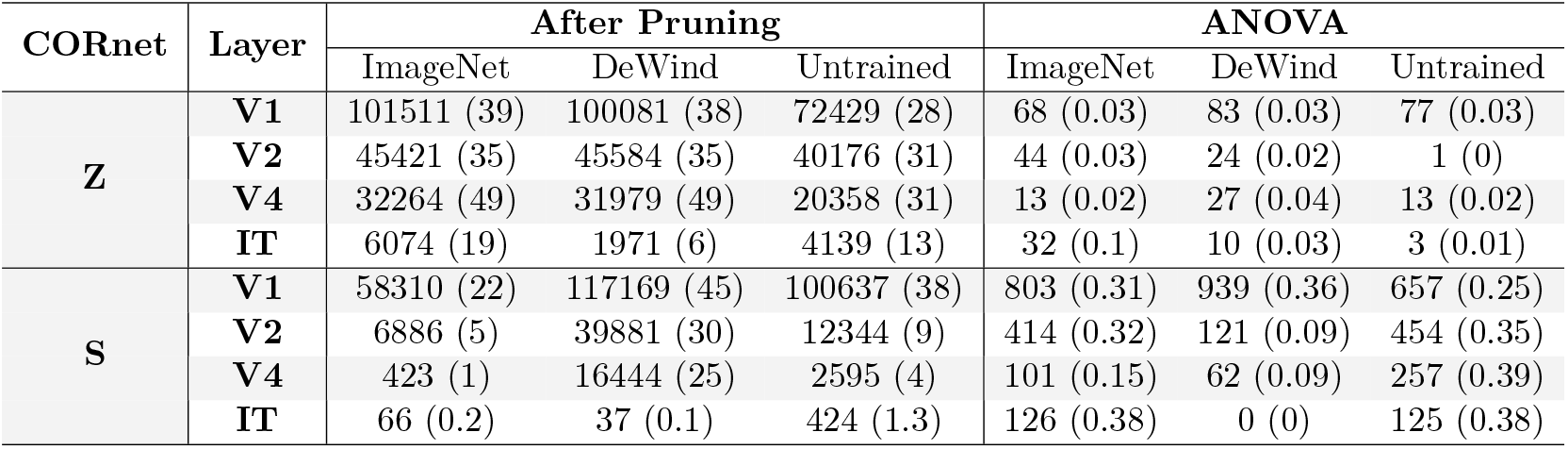
Number of retained units after pruning, and of number-detector units identified by ANOVA. The full set of units in V1, V2, V4, and IT layer in both models are 262144, 131072, 65536, 32768 respectively. The numbers in parentheses denote the percentage of units compared to the full set.

### 3.2 Retained Units after Pruning Fit the Behavior Data Better than Number-Detector Units

Figure 2 shows that the retained units after pruning provide a better fit for modeling the number RDM compared to the full set of units, highlighting the limitations of using the full set in classical RSA. Additionally, in most cases—except for V2 and V1 in CORnet-S DeWind, which is specifically trained for number discrimination—the number-detector units selected by ANOVA, as commonly studied in traditional literature, perform worse than the retained units after pruning. In 8/22 cases, they are even worse than the full set of units. This demonstrates that the number-detector units are not important for capturing the population-level representation of numerosity in human behavioral data.

**Figure 2.**
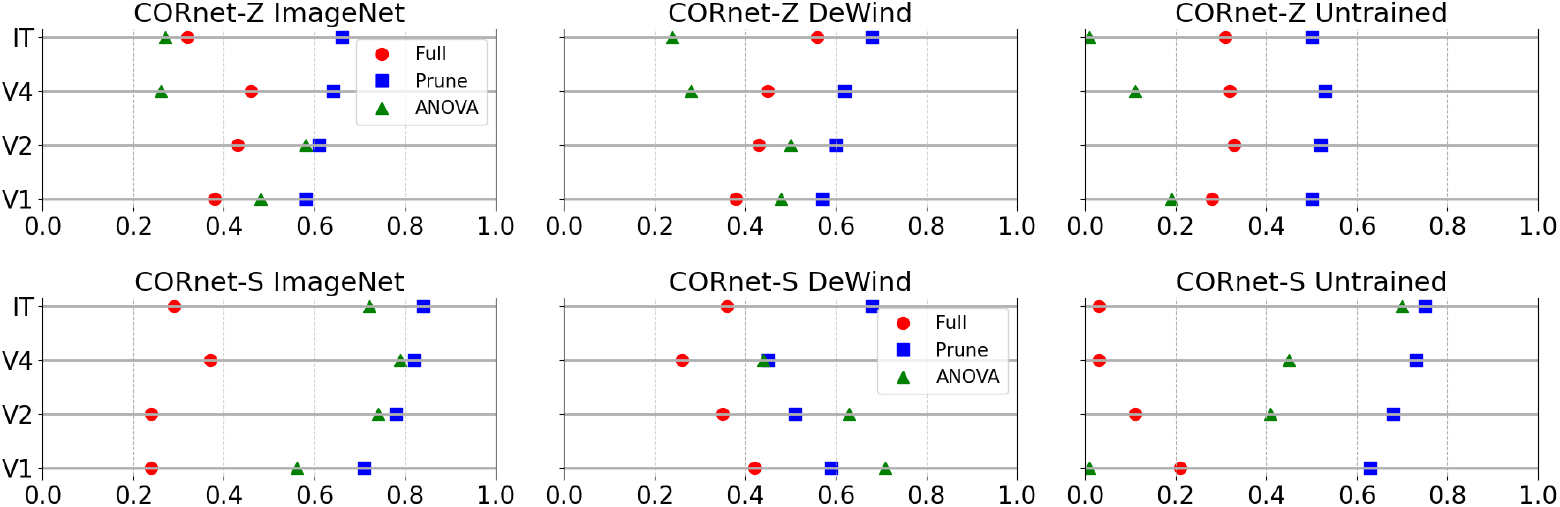
The Pearson correlations, quantified using RSA, from the full set of units, the retained units after pruning, and the number-detector units identified by ANOVA. The missing data points are due to an insufficient number of minimum units (2) required to compute the correlations.

## 4 Discussion

It has been emphasized that cognitive and behavioral functions emerge from the collective dynamics of neural populations, rather than isolated neuronal activity (Yuste 2015), underscoring the importance of population-level analysis for understanding complex behaviors like numerosity discrimination.

In this context, RSA has become a widely used method for comparing representational spaces from human brain activity, behavioral data, and computational models. However, classical RSA assumes that each feature is equally important, making it difficult to interpret the contribution of individual features. In our case, classical RSA does not provide insights into the importance of number-detector units identified in CNNs. To address this limitation, we applied pruning, a feature selection technique used to identify and retain the most relevant components of a model—such as specific weights or activations—that best align with behavioral data. Pruning revealed that almost none of the number-detector units contribute to the representation of numerosity at the population level, casting doubt on their significance for numerosity discrimination. These findings are also consistent with previous work by Mistry et al. (2023), which demonstrated that training a CNN for numerosity discrimination significantly reorganizes the number-detector units, and by Chapalain, Thirion, and Eger (2024), which showed that dot-pattern-tuned units do not generalize to object-number information in photorealistic stimuli.

## Acknowledgments

This paper was previously presented at the NeurIPS 2024 Workshop on Behavioral Machine Learning. More details about the workshop are available at [NeurIPS 2024 Behavioral Machine Learning Workshop](https://sites.google.com/view/behavioralml/).

## Declaration of Generative AI and AI-assisted Technologies

The authors used ChatGPT to assist with rephrasing certain sentences during the preparation of this work. They subsequently reviewed and edited the content as needed and took full responsibility for the final published article.

## Data and Code Availability

All code and data used to generate stimuli, train models, and analyze results is available at: https://github.com/alireza-kr/CORnetOnNumber

## Notes

https://github.com/alireza-kr/CORNumUnit

### Competing Interest Statement

The authors have declared no competing interest.

## References

Bao, Wanqian and Uri Hasson (2024). “Identifying and interpreting non-aligned human conceptual representations using language modeling”. In: arXiv preprint arXiv:2403.06204.

Castaldi, Elisa et al.. (July 2019). “Attentional amplification of neural codes for number independent of other quantities along the dorsal visual stream”. In: eLife 8. doi: 10.7554/elife.45160. url: https://doi.org/10.7554/elife.45160.

Chapalain, Thomas, Bertrand Thirion, and Evelyn Eger (Sept. 2024). “Trained deep neural network models of the ventral visual pathway encode numerosity with robustness to object and scene identity”. In: bioRxiv. doi: 10.1101/2024.09.05.611433. url: https://doi.org/10.1101/2024.09.05.611433.

Cheng, Yu et al.. (2015). “An exploration of parameter redundancy in deep networks with circulant projections”. In: Proceedings of the IEEE international conference on computer vision, pp. 2857–2865.

Cichy, Radoslaw Martin et al.. (June 2016). “Comparison of deep neural networks to spatio-temporal cortical dynamics of human visual object recognition reveals hierarchical correspondence”. In: Scientific Reports 6.1. doi: 10.1038/srep27755. url: https://doi.org/10.1038/srep27755.

Dehaene, Stanislas (Apr. 2011). The number sense. OUP USA.

Deng, Jia et al.. (June 2009). “ImageNet: A large-scale hierarchical image database”. In: 2009 IEEE Conference on Computer Vision and Pattern Recognition. doi: 10.1109/cvpr.2009.5206848. url: https://doi.org/10.1109/cvpr.2009.5206848.

DeWind, Nicholas K. et al.. (Sept. 2015). “Modeling the approximate number system to quantify the contribution of visual stimulus features”. In: Cognition 142, pp. 247–265. doi: 10.1016/j.cognition.2015.05.016. url: https://doi.org/10.1016/j.cognition.2015.05.016.

Edwards, Laura A. et al.. (Sept. 2015). “Functional brain organization for number processing in pre-verbal infants”. In: Developmental Science 19.5, pp. 757–769. doi: 10.1111/desc.12333. url: https://doi.org/10.1111/desc.12333.

Flechas Manrique, Natalia et al. (Dec. 2023). “Enhancing Interpretability Using Human Similarity Judgements to Prune Word Embeddings”. In: Proceedings of the 6th BlackboxNLP Workshop: Analyzing and Interpreting Neural Networks for NLP. Ed. by Yonatan Belinkov et al. Singapore: Association for Computational Linguistics, pp. 169–179. doi: 10.18653/v1/2023.blackboxnlp-1.13. url: https://aclanthology.org/2023.blackboxnlp-1.13.

Frankle, Jonathan and Michael Carbin (2018). “The lottery ticket hypothesis: Finding sparse, trainable neural networks”. In: arXiv preprint arXiv:1803.03635.

Hyde, Daniel C. et al.. (Nov. 2010). “Near-infrared spectroscopy shows right parietal specialization for number in pre-verbal infants”. In: NeuroImage 53.2, pp. 647–652. doi: 10.1016/j.neuroimage.2010.06.030. url: https://doi.org/10.1016/j.neuroimage.2010.06.030.

Izard, Véronique, Ghislaine Dehaene-Lambertz, and Stanislas Dehaene (Feb. 2008). “Distinct cerebral pathways for object identity and number in human infants”. In: PLoS Biology 6.2, e11. doi: 10.1371/journal.pbio.0060011. url: https://doi.org/10.1371/journal.pbio.0060011.

Kaniuth, Philipp and Martin N. Hebart (Aug. 2022). “Feature-reweighted representational similarity analysis: A method for improving the fit between computational models, brains, and behavior”. In: NeuroImage 257, p. 119294. doi: 10.1016/j.neuroimage.2022.119294. url: https://doi.org/10.1016/j.neuroimage.2022.119294.

Karami, Alireza (2024). “The representation of numerosity in the human brain and machines”. Available at: https://doi.org/10.15168/11572_402591. PhD thesis. doi: 10.15168/11572_402591.

Karami, Alireza, Elisa Castaldi, Evelyn Eger, Martin Hebart, et al. (Nov. 16, 2025). “Numerosity Is Directly Sensed and Dynamically Transformed in the Human Brain: Evidence from MEG-MRI Fusion”. In: bioRxiv. doi: 10.1101/2025.11.15.687894. url: https://doi.org/10.1101/2025.11.15.687894.

Karami, Alireza, Elisa Castaldi, Evelyn Eger, and Manuela Piazza (July 9, 2025). “Distinct neural representational geometries of numerosity in early visual and association regions across visual streams”. In: Communications Biology 8.1, p. 1029. doi: 10.1038/s42003-025-08395-z. url: https://doi.org/10.1038/s42003-025-08395-z.

Karami, Alireza, Nhut Truong, and Manuela Piazza (Jan. 1, 2025). “Investigation of Numerosity Representation in Convolution Neural Networks”. In: CCN 2025 Proceedings. doi: 10.32470/4j01408. url: https://doi.org/10.32470/4j01408.

Khaligh-Razavi, Seyed-Mahdi and Nikolaus Kriegeskorte (Nov. 2014). “Deep supervised, but not unsupervised, models may explain IT cortical representation”. In: PLoS Computational Biology 10.11, e1003915. doi: 10.1371/journal.pcbi.1003915. url: https://doi.org/10.1371/journal.pcbi.1003915.

Kim, Gwangsu et al.. (Jan. 2021). “Visual number sense in untrained deep neural networks”. In: Science Advances 7.1. doi: 10.1126/sciadv.abd6127. url: https://doi.org/10.1126/sciadv.abd6127.

Kobylkov, Dmitry et al.. (Aug. 2022). “Number neurons in the nidopallium of young domestic chicks”. In: Proceedings of the National Academy of Sciences 119.32. doi: 10.1073/pnas.2201039119. url: https://doi.org/10.1073/pnas.2201039119.

Kriegeskorte, Nikolaus (Jan. 2008). “Representational similarity analysis – connecting the branches of systems neuroscience”. In: Frontiers in Systems Neuroscience. doi: 10.3389/neuro.06.004.2008. url: https://doi.org/10.3389/neuro.06.004.2008.

Kruskal, J. B. (Mar. 1964). “Multidimensional scaling by optimizing goodness of fit to a nonmetric hypothesis”. In: Psychometrika 29.1, pp. 1–27. doi: 10.1007/bf02289565. url: https://doi.org/10.1007/bf02289565.

Kubilius, Jonas, Martin Schrimpf, Kohitij Kar, et al. (Oct. 2019). “Brain-Like Object Recognition with High-Performing Shallow Recurrent ANNs”. In: arXiv.

Kubilius, Jonas, Martin Schrimpf, Aran Nayebi, et al. (Sept. 2018). “CORNET: Modeling the neural mechanisms of core object recognition”. In: bioRxiv (Cold Spring Harbor Laboratory). doi: 10.1101/408385. url: https://doi.org/10.1101/408385.

Lasne, Gabriel et al.. (May 2019). “Discriminability of numerosity-evoked fMRI activity patterns in human intra-parietal cortex reflects behavioral numerical acuity”. In: Cortex 114, pp. 90–101. doi: 10.1016/j.cortex.2018.03.008. url: https://doi.org/10.1016/j.cortex.2018.03.008.

Lindsay, Grace W. (Sept. 2021). “Convolutional neural networks as a model of the visual system: past, present, and future”. In: Journal of Cognitive Neuroscience 33.10, pp. 2017–2031. doi: 10.1162/jocn\{_}a\{_}01544. url: https://doi.org/10.1162/jocn_a_01544.

Mistry, Percy K. et al.. (June 2023). “Learning-induced reorganization of number neurons and emergence of numerical representations in a biologically inspired neural network”. In: Nature Communications 14.1. doi: 10.1038/s41467-023-39548-5. url: https://doi.org/10.1038/s41467-023-39548-5.

Nasr, Khaled, Pooja Viswanathan, and Andreas Nieder (May 2019). “Number detectors spontaneously emerge in a deep neural network designed for visual object recognition”. In: Science Advances 5.5. doi: 10.1126/sciadv.aav7903. url: https://doi.org/10.1126/sciadv.aav7903.

Park, Joonkoo (Dec. 2022). “Flawed stimulus design in additive-area heuristic studies”. In: Cognition 229, p. 104919. doi: 10.1016/j.cognition.2021.104919. url: https://doi.org/10.1016/j.cognition.2021.104919.

Piazza, Manuela et al.. (Oct. 2004). “Tuning curves for approximate numerosity in the human intraparietal sulcus”. In: Neuron 44.3, pp. 547–555. doi: 10.1016/j.neuron.2004.10.014. url: https://doi.org/10.1016/j.neuron.2004.10.014.

Schrimpf, Martin et al.. (Sept. 2018). “Brain-Score: Which Artificial Neural Network for Object Recognition is most Brain-Like?” In: bioRxiv (Cold Spring Harbor Laboratory). doi: 10.1101/407007. url: https://doi.org/10.1101/407007.

Tarigopula, Priya et al.. (Nov. 2023). “Improved prediction of behavioral and neural similarity spaces using pruned DNNs”. In: Neural Networks 168, pp. 89–104. doi: 10.1016/j.neunet.2023.08.049. url: https://doi.org/10.1016/j.neunet.2023.08.049.

Truong, Nhut, Dario Pesenti, and Uri Hasson (2024). “Explaining Human Comparisons using Alignment-Importance Heatmaps”. In: Proceedings of the Annual Meeting of the Cognitive Science Society. Vol. 46.

Viswanathan, Pooja and Andreas Nieder (June 2013). “Neuronal correlates of a visual “sense of number” in primate parietal and prefrontal cortices”. In: Proceedings of the National Academy of Sciences 110.27, pp. 11187–11192. doi: 10.1073/pnas.1308141110. url: https://doi.org/10.1073/pnas.1308141110.

Wagener, Lysann et al.. (Apr. 2018). “Neurons in the endbrain of numerically naive crows spontaneously encode visual numerosity”. In: Current Biology 28.7, 1090–1094.e4. doi: 10.1016/j.cub.2018.02.023. url: https://doi.org/10.1016/j.cub.2018.02.023.

Yamins, Daniel L K and James J DiCarlo (Feb. 2016). “Using goal-driven deep learning models to understand sensory cortex”. In: Nature Neuroscience 19.3, pp. 356–365. doi: 10.1038/nn.4244. url: https://doi.org/10.1038/nn.4244.

Yamins, Daniel L. K. et al.. (May 2014). “Performance-optimized hierarchical models predict neural responses in higher visual cortex”. In: Proceedings of the National Academy of Sciences 111.23, pp. 8619–8624. doi: 10.1073/pnas.1403112111. url: https://doi.org/10.1073/pnas.1403112111.

Yuste, Rafael (July 2015). “From the neuron doctrine to neural networks”. In: Nature reviews. Neuroscience 16.8, pp. 487–497. doi: 10.1038/nrn3962. url: https://doi.org/10.1038/nrn3962.

Zhang, Xi and Xiaolin Wu (Nov. 2020). On numerosity of deep neural networks. url: https://arxiv.org/abs/2011.08674.

